# Orphan nuclear receptors recruit TRIM28 to promote telomeric H3K9me3 for the alternative lengthening of telomeres pathway

**DOI:** 10.1101/2025.06.16.658187

**Authors:** Chia-Tsen Tsai, Venus Marie Gaela, Hsuan-Yu Hsia, Yu-Chen Huang, Liuh-Yow Chen

## Abstract

Alternative lengthening of telomeres (ALT) is a telomere maintenance mechanism deployed in embryonic stem cells and cancer cells. High levels of the heterochromatin mark H3 lysine 9 trimethylation (H3K9me3) at telomeres are critical for ALT, but how that is achieved remains unclear. Telomeric association of orphan nuclear receptors (NRs)—such as COUP-TF1, COUP-TF2, TR2, and TR4—has been shown previously to promote ALT activation. Here, we show that orphan NRs regulate telomeric H3K9me3 through TRIM28, a corepressor of ZNF transcription factors, to drive ALT. We report that H3K9me3 is induced by telomeric association of orphan NRs in cultured human fibroblast and ALT cancer cell lines. Moreover, TRIM28 is required for the orphan NR-induced H3K9me3 and ALT phenotypes. Importantly, physical interaction of TRIM28 with orphan NRs facilitates a telomeric localization of TRIM28. A TRIM28 variant defective in orphan NR interaction fails to localize to telomeres and is unable to promote H3K9me3 and ALT phenotypes. These findings indicate that telomeric orphan NRs recruit TRIM28, driving telomeric H3K9me3 and ALT activation, emphasizing the role of changes in chromatin structure in ALT activation.

## Introduction

Telomeres, composed of repetitive DNA sequences at chromosome ends, are crucial for genomic stability by preventing their recognition as double-strand breaks. This protective mechanism is mediated by the telomere-associated shelterin protein complex, which safeguards telomeres from initiating DNA damage response (Moyzis et al., 1988; Palm & De Lange, 2008). In somatic cells, cell division results in telomere shortening due to the end-replication problem, ultimately leading to cellular senescence(Harley et al., 1990; Wright & Shay, 1992). Cancer cells overcome this limitation by adopting telomere maintenance mechanisms, either by reactivating telomerase or activating the alternative lengthening of telomeres (ALT) pathway. ALT enacts homologous recombination (HR), reminiscent of break-induced replication, to lengthen telomeres and enable indefinite proliferation(Bryan et al., 1997; Dilley et al., 2016; Zhang et al., 2019). It is characterized by several molecular hallmarks. For instance, in ALT cells, telomeres are encapsulated in promyelocytic leukemia nuclear bodies (PML-NBs), forming ALT-associated PML-NBs (APBs)(Yeager et al., 1999). APBs are hubs where telomere DNA and DNA damage response factors cluster, together catalyzing ALT recombination and break-induced replication(Cho et al., 2014; Zhang et al., 2021). In addition, telomeric DNA in ALT cells exhibits spontaneous damage due to replication stress, resulting in telomere excision and accumulation of extrachromosomal telomeric repeats (ECTRs) (Cesare et al., 2009; Cesare & Griffith, 2004; Chen et al., 2017; Wang et al., 2004). Moreover, levels of telomeric repeat-containing RNAs (TERRAs), which can form RNA:DNA hybrids that are sources of replication stress, are elevated in ALT cells (Chu et al., 2017; Feretzaki et al., 2020). Thus, these ALT phenotypes reflect ALT activation and influence oncogenesis.

Regulation of telomere chromatin plays a critical role in ALT activation. Zinc finger protein 827 (ZNF827) recruits the nucleosome remodeling and histone deacetylation (NuRD) complex to ALT telomeres, leading to histone deacetylation (Conomos et al., 2014). In turn, histone deacetylation depletes shelterin from telomeres and promotes telomere interaction and compaction. Additionally, the ZNF827-NuRD complex recruits HR-associated proteins to telomeres and, through its chromatin remodeling function, promotes ALT activity(Conomos et al., 2014). Involvement of the ZNF827-NuRD complex in ALT telomere maintenance may be related to the role of the NuRD complex in heterochromatin assembly during DNA replication(Sims & Wade, 2011).

Previous studies have shown that ALT telomeric chromatin is less compact yet highly heterochromatic compared to telomerase-positive telomeric chromatin(Episkopou et al., 2014). The heterochromatin mark histone H3 lysine 9 trimethylation (H3K9me3) is enriched at ALT telomeres, and its levels affect ALT activity. Telomeres in mouse embryonic stem cells (mESCs) exhibit heterochromatic features, ALT-like phenotypes, and increased recombination. A proteomic study of telomere chromatin from mESCs revealed the telomeric association of the SET Domain Bifurcated 1 (SETDB1) histone methyltransferase, which is required for telomeric H3K9me3 and also regulates APB formation in mESCs(Gauchier et al., 2019). SETDB1 is also associated with ALT telomeres in cancer cells, where its depletion reduces telomeric H3K9me3 and ALT phenotypes, supporting the functional significance of heterochromatin to ALT activation(Gauchier et al., 2019). Interestingly, SETDB1 also regulates TERRA transcription, indicating that telomeric heterochromatin is compatible with transcriptional elongation at telomeres (Gauchier et al., 2019). The telomeric localization and protein stability of SETDB1 in ALT cells both depend on Tripartite motif-containing 28 (TRIM28), a corepressor of the family of Krüppel-associated box zinc finger proteins (KRAB-ZFP) (Friedman et al., 1996; Moosmann et al., 1996; Schultz et al., 2001). Consistent with its SETDB1 interaction, TRIM28 promotes occupancy of H3K9me3 on telomeres in ALT cells(Wang et al., 2021). However, depletion of TRIM28 results in telomere lengthening and inconsistent ALT phenotypes. This paradox underscores the need for further research to clarify the role of heterochromatin and TRIM28 in ALT telomere maintenance.

Moreover, ALT activation is associated with a deficiency in the histone H3.3 chaperone complex α-thalassemia/mental retardation syndrome X-linked (ATRX) and death-domain-associated protein (DAXX). Restoration of ATRX expression in ATRX-negative ALT cell lines has been shown to suppress many ALT-associated phenotypes(Clynes et al., 2015; Napier et al., 2015). However, knockdown of ATRX or DAXX in either mortal or telomerase-positive cell lines results in reduced histone deposition and telomeric H3K9me3, telomere decompaction, and replication dysfunction, though loss of either protein is not sufficient to immediately activate ALT(Li et al., 2019; Lovejoy et al., 2012; O’Sullivan et al., 2014). Intriguingly, Gauchier et al. demonstrated that ATRX depletion induces ALT recombination in mESCs that is dependent on H3K9me3(Gauchier et al., 2019), indicating that ALT activation may require the combined effects of ATRX deficiency and increased levels of H3K9me3 at telomeres. However, the mechanism driving formation of telomeric H3K9me3 in ALT cells remains to be determined.

Orphan nuclear receptors (NRs) of the NR2C/F class—such as COUP-TF1, COUP-TF2, TR2, TR4, and EAR2—associate with telomeres in ALT cells by binding to TCAGGG telomere variant repeats and they regulate ALT activity (Déjardin and Kingston, 2009; Conomos et al., 2012). These orphan NRs facilitate recruitment of the ZNF827-NuRD complex to telomeres, which remodels telomeric chromatin and enhances recombination (Conomos, et al., 2014). Additionally, orphan NRs bind directly to FANCD2, a crucial component of the Fanconi anemia repair pathway, to initiate a DNA damage response that boosts ALT activity in ALT cells (Xu et al., 2019). Recently, we developed a system to ectopically express orphan NRs, tethering them to telomeres in non-ALT fibroblasts to explore the molecular mechanisms by which orphan NRs regulate ALT(Gaela et al., 2024). Our previous findings indicate that the telomeric association of orphan NRs is sufficient to trigger APB formation and induce various ALT phenotypes and telomere recombination. This ALT induction is mediated by activation function 2 (AF2) domains of orphan NRs and ZNF827, indicating an involvement of altered chromatin structures in orphan NR-induced ALT activation. These findings highlight the critical role of orphan NRs in ALT regulation (Gaela et al., 2024). Building on this system, in the current study, we aimed to investigate how orphan NRs contribute to ALT chromatin regulation through TRIM28, with a particular focus on elucidating the mechanisms by which H3K9me3-marked heterochromatin is established at telomeres for ALT activation.

## Result

### Orphan NRs promote telomeric H3K9me3

The telomeric association of orphan NRs has been shown previously to promote ALT(Conomos et al., 2012, 2014), yet the exact mechanism remains to be established. Structural alterations in telomeric chromatin, including changes in histone modifications such as H3K9me3 and H4 acetylation (H4ac), have been linked to ALT activation (Conomos et al., 2014; Gauchier et al., 2019). To determine how orphan NRs regulate the telomeric epigenetic structures of ALT cells, first, we performed telomere-chromatin immunoprecipitation (telomere-ChIP) to detect histone H3K9me3 and H4ac in cultured ALT cell lines, including U2OS osteosarcoma cells and transformed WI38-VA13/2RA fibroblasts. Our results revealed that both these histone modifications are present on telomeric histones (Figure 1B). Given that COUP-TF2 and TR4 are highly expressed in U2OS and WI38-VA13/2RA cells and are known to localize to telomeres(Conomos et al., 2012, 2014; Gaela et al., 2024), we silenced their expression by means of small interfering RNAs (siRNAs) (Figure 1A) to assess their impact on telomeric H3K9me3 and H4ac. We observed that simultaneous knockdown of COUP-TF2 and TR4 from U2OS and WI38-VA13/2RA cells caused a significant decrease in the levels of telomeric H3K9me3, but not in H4ac levels (Figure 1B), indicating that orphan NRs specifically modulate H3K9me3 at telomeres in ALT cells.

**Figure 1.**
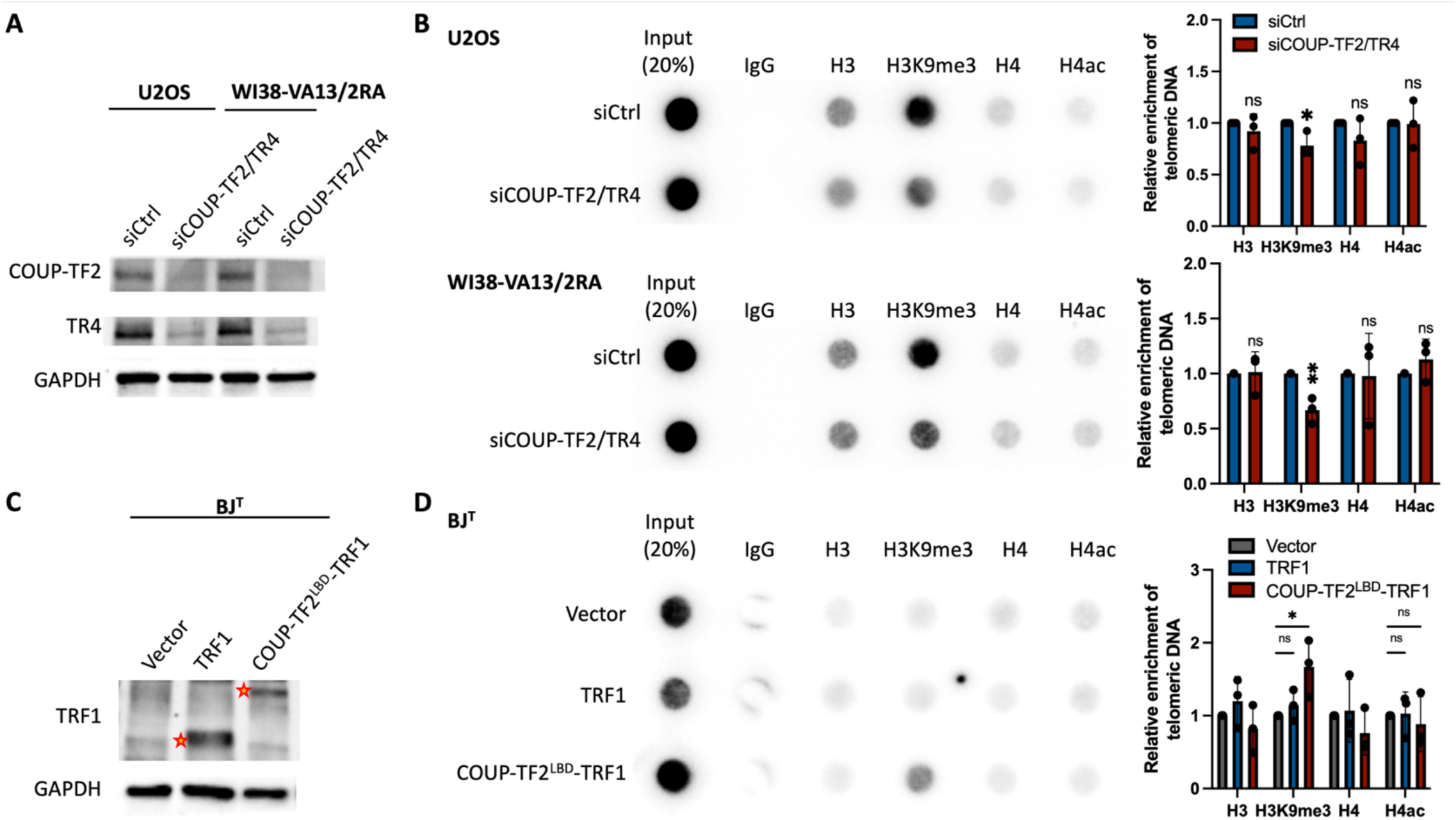
Role of orphan nuclear receptors (NRs) in enhancing telomeric H3K9me3 levels. (**A**) Western blot analysis showing the expression levels of COUP-TF2 and TR4 in U2OS and WI38-VA13/2RA cells following treatment for 6 days with specific siRNAs targeting these NRs. (**B**) Telomere-ChIP analysis of U2OS and WI38-VA13/2RA cells 6 days after COUP-TF2 and TR4 knockdown to assess enrichment for telomeric DNA with the indicated histones and histone modifications (H3=histone 3, H3K9me3=histone H3 lysine 9 trimethylation, H4=histone 4, H4ac=pan histone H4 acetylation). The relative enrichment was calculated after normalization of ChIP DNA signals to the respective input DNA signals. Results from three independent experiments are shown as individual data points (mean ± SD; n = 3 independent experiments). (**C**) Western blot analysis of BJ^T^ cells ectopically overexpressing TRF1 alone or COUP-TF2^LBD^-TRF1, confirming successful expression of the constructs with an anti-TRF1 antibody. (D) Telomere-ChIP analysis on BJ^T^ cells overexpressing TRF1 alone or COUP-TF2^LBD^-TRF1, showing enrichment for telomeric DNA with the indicated histones and histone marks. The relative enrichment was calculated after normalization of ChIP DNA signals to the respective input DNA signals. Results from three independent experiments are shown as individual data points. (mean ± SD; n = 3 independent experiments). (B, D) are determined by the unpaired t-test. Statistical significance is noted as follows: ns (P > 0.05), * (P < 0.05), ** (P < 0.01).

We have previously demonstrated that targeting orphan NRs to telomeres is sufficient to induce ALT activity and various ALT features in primary human fibroblast cells(Gaela et al., 2024). Adopting the same strategy herein, we endeavored to uncover a direct role for orphan NRs in regulating telomere chromatin. Consistently, we found that ectopic expression of the COUP-TF2 ligand-binding domain (LBD) fused with TRF1 (COUP-TF2^LBD^-TRF1), but not TRF1 alone, in telomerase-immortalized human BJ fibroblast cells (BJ^T^) led to ALT induction, which was characterized by co-localization of telomeres with PML-NBs, thereby forming APBs, and by the occurrence of non-S phase ALT telomere DNA synthesis (ATDS) (Figure 1C, Supplementary Figure 1A, B). Furthermore, telomere-ChIP revealed an increase in telomeric H3K9me3 modification, but not of H4ac, in BJ^T^ cells expressing COUP-TF2^LBD^-TRF1 relative to Vector (Figure 1D). This result indicates that telomeric targeting of COUP-TF2 LBD can promote histone H3K9me3 on telomeres. Collectively, these data reveal that the telomere association of orphan NRs in ALT cells promotes structural changes in telomeric chromatin by inducing telomeric H3K9me3 occupancy.

### Alteration of telomere chromatin structures promotes ALT activity and associated features

As evidenced above, structural alterations in telomeric chromatin are associated with orphan NR-mediated ALT induction, consistent with a link between distinct telomeric chromatin environments and ALT maintenance(Li et al., 2019; O’Sullivan et al., 2014; Wang et al., 2021). To further investigate the causative role of chromatin states in ALT induction, we ectopically expressed in BJ^T^ cells the transcriptional repressor Krüppel associated box (KRAB) or the transcriptional activator herpes simplex virus VP64, both of which were fused to TRF1. These fusion proteins were used to specifically induce the formation of heterochromatin and euchromatin, respectively, at the telomeres of the BJ^T^ cells. First, we confirmed transcriptional effector activities by assessing expression of TERRAs by means of reverse transcription polymerase chain reaction (RT-PCR)(Feretzaki & Lingner, 2017). As anticipated, we observed that TERRA expression was suppressed by KRAB-TRF1 and upregulated by VP64-TRF1 relative to cells expressing TRF1 alone (Figure 2A, Supplementary Figure 2A). Furthermore, we assessed the effect of telomeric targeting of KRAB or VP64 in BJ^T^ cells on chromatin modifications through telomere-ChIP. Our results show that KRAB-TRF1 significantly increased telomeric histone occupancies by H3, H4, and H3K9me3, indicative of telomere compaction. In contrast, VP64-TRF1 expression reduced telomere histone occupancy and H3K9me3 levels (Figure 2B). Together, these results demonstrate that KRAB-TRF1 and VP64-TRF1 induce heterochromatin and euchromatin formation, respectively, at telomeres in BJ^T^ cells. Next, we examined ALT induction following these chromatin alterations. Robust APB formation and ATDS were observed in the BJ^T^ cells expressing KRAB-TRF1, whereas VP64-TRF1 expression only induced minimal APB formation and ATDS (Figure 2C, D). Notably, these ALT features were absent from cells expressing empty vector, TRF1 alone, KRAB alone, or VP64 alone. Together, these findings support that alterations in chromatin structure may drive ALT induction.

**Figure 2.**
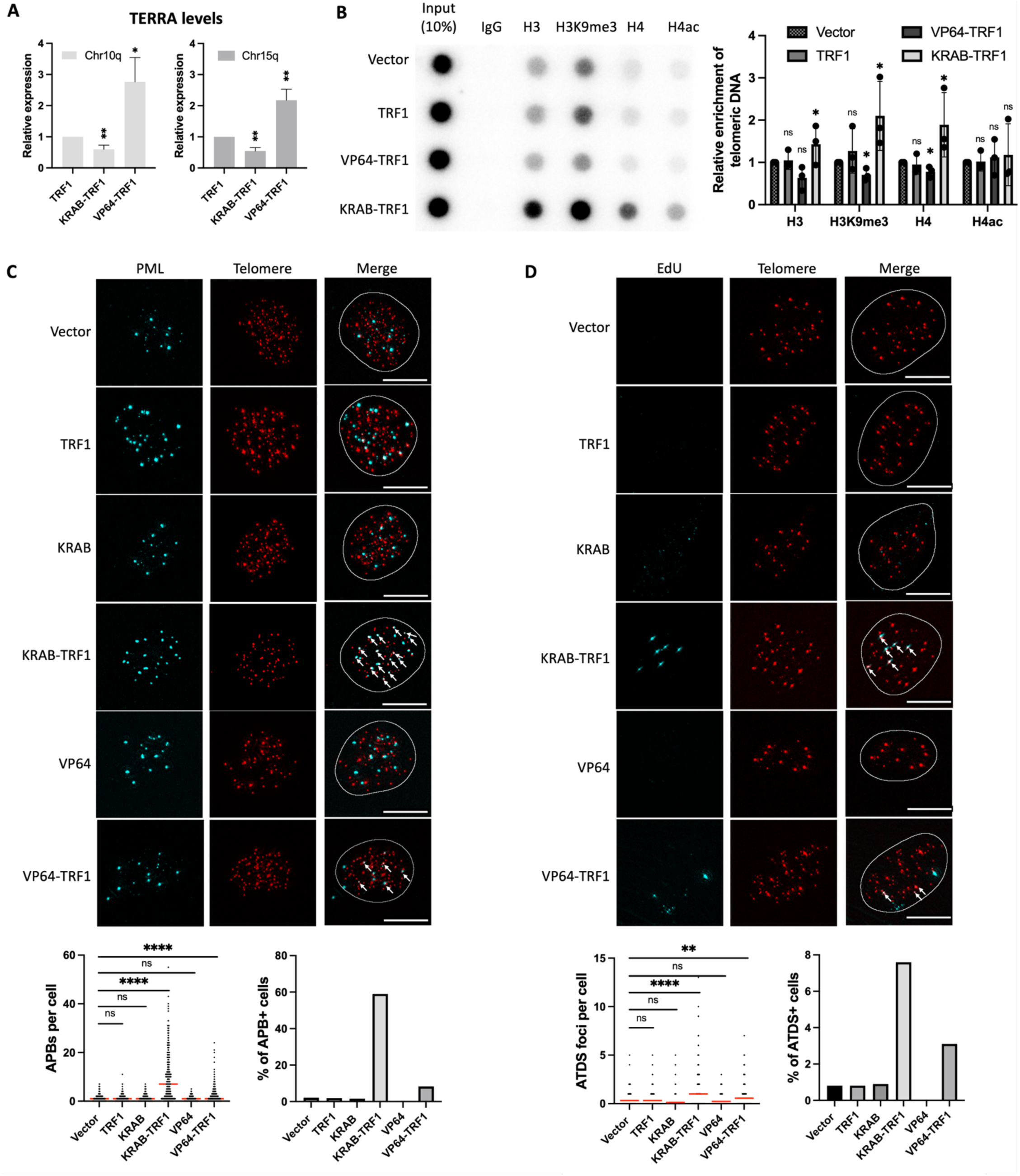
KRAB-TRF1-mediated heterochromatin formation enhances ALT induction. (**A**) Quantification of TERRA expression levels in KRAB-TRF1 or VP64-TRF1 BJ^T^ cells, normalized to those of TRF1-expressing BJ^T^ cells, as determined by qPCR of RNA samples. TERRA levels for different subtelomeric sequences from the chromosome (Chr) 10q or 15q ends were measured and normalized to GAPDH (mean ± SEM; n = 5 independent experiments). (**B**) Telomere-ChIP analysis in BJ^T^ cells overexpressing TRF1, KRAB-TRF1, or VP64-TRF1 reveals enrichment for telomeric DNA with indicated histones and histone modifications (H3=histone 3, H3K9me3=histone H3 lysine 9 trimethylation, H4=histone 4, H4ac=pan histone H4 acetylation). The relative enrichment was calculated after normalization of ChIP DNA signals to the respective input DNA signals. Results from three independent experiments are shown as individual data points (mean ± SD; n = 3 independent experiments). (**C**) Representative immunofluorescence images depicting the localization of PML and telomeres as APBs in BJ^T^ cells expressing different plasmids. Immunofluorescence (IF) identifies ectopically expressed proteins and PML, whereas telomeres are visualized by FISH using the TelC PNA probe. Co-localization (white arrows) of PML (cyan) and telomeres (red) appears white. White outlines encompass DAPI segmentation. The dot plot shows the quantitative numbers of APBs in individual BJ^T^ cells (n > 150). A bar chart illustrates the percentages of APB-positive (APB+) cells, defined as those containing more than five APBs. (**D**) Representative images showing EdU incorporation at telomeres in BJ^T^ cells detected by Click chemistry. Telomeres are visualized by FISH using the TelC PNA probe. Co-localization (white arrows) of EdU (cyan) and telomeres (red) appears white. White outlines encompass DAPI segmentation. The dot plot shows the quantitative numbers of the co-localization of telomeres with EdU in BJ^T^ cells (n > 150). A bar chart displays the percentage of ATDS-positive (ATDS+) cells containing more than three EdU+Telomere foci. Red lines indicate the mean. (A, B) are determined by the unpaired t-test. (C, D) are determined by the Mann-Whitney U test. Statistical significance is denoted as follows: ns (P > 0.05), * (P < 0.05), ** (P < 0.01), *** (P < 0.001), **** (P < 0.0001), as determined by the Mann-Whitney U test.

### TRIM28 promotes telomeric H3K9me3 and ALT induction

When associated with telomeres, orphan NRs and KRAB promote H3K9me3 and trigger ALT induction, consistent with the notion that changes in chromatin structure may underlie ALT activation. To further elucidate the significance of chromatin changes for orphan NR-mediated ALT induction, we silenced the expression of KRAB-associated chromatin regulators, such as SET domain bifurcated histone lysine methyltransferase 1 (SETDB1), Heterochromatin Protein 1 (HP1)α/β/γ, Histone deacetylase (HDAC) 1/2 or DNA (cytosine-5)-methyltransferase 1 (DNMT1) in COUP-TF2^LBD^-TRF1 BJ^T^ cells(Czerwińska et al., 2017)(Supplementary Figure. 3A, B). We found that depletion of any of these chromatin regulators reduced APB formation and ATDS (Supplementary Figure 3C, D), supporting that chromatin alterations may contribute to orphan NR-induced ALT activation.

Intriguingly, SETDB1, HP1α/β/γ, HDAC1/2, and DNMT1 have all been shown previously to associate with TRIM28 and are recruited together for gene regulation by KRAB domain-containing zinc-finger transcription factors via TRIM28-KRAB domain interactions (Friedman et al., 1996). Moreover, Wang et al. recently reported that TRIM28 localizes to telomeres in ALT cells and impacts H3K9me3 (2021), suggesting that TRIM28 might be involved in orphan NR-mediated ALT regulation. To examine this possibility, we silenced TRIM28 expression in BJ^T^ cells by means of siRNAs to investigate its effects on COUP-TF2^LBD^-TRF1-mediated telomeric H3K9me3 and ALT induction (Figure 3A). Telomere-ChIP experiments revealed that TRIM28 depletion had a limited effect on telomeric H3K9me3 in vector-administered BJ^T^ cells, whereas levels of H3K9me3 in the COUP-TF2^LBD^-TRF1-expressing cells were significantly reduced (Figure 3B). These results indicate that TRIM28 regulates COUP-TF2^LBD^-induced telomeric H3K9me3, but not maintenance of basal telomeric H3K9me3 in BJ^T^ cells (Figure 3B). Furthermore, knockdown of TRIM28 reduced APB formation and ATDS in BJ^T^ cells expressing COUP-TF2^LBD^-TRF1 (Figure 3C-E). Together, these findings support that TRIM28 plays a crucial role in COUP-TF2-induced ALT induction by regulating telomere chromatin in human fibroblasts.

**Figure 3.**
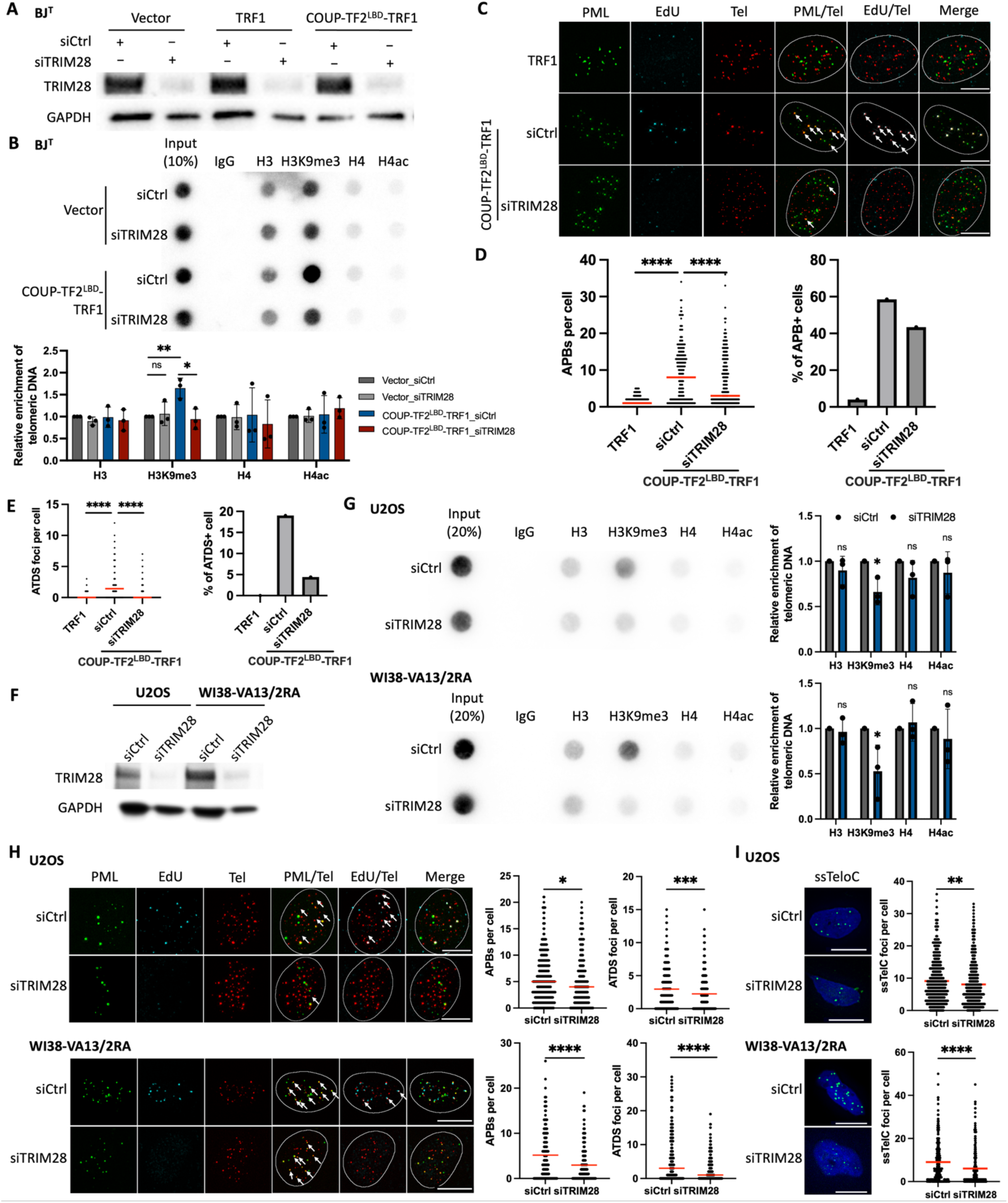
TRIM28 mediates the enrichment of H3K9m3 and ALT activation. (**A**) Western blot analysis of TRIM28 expression in BJ^T^ cells after 6 days of treatment with TRIM28-targeting siRNAs. (**B**) Telomere-ChIP analysis of BJ^T^ cells 6 days following TRIM28 knockdown to assess enrichment for telomeric DNA with indicated histones and histone modifications (H3=histone 3, H3K9me3=histone H3 lysine 9 trimethylation, H4=histone 4, H4ac=pan histone H4 acetylation). The relative enrichment was calculated after normalization of ChIP DNA signals to the respective input DNA signals (mean ± SD; n = 3 independent experiments). Results from three independent experiments are shown as individual data points. (**C**) Representative images showing EdU incorporation at telomeres in BJ^T^ cells detected by Click chemistry. PML was detected by IF, and telomeres were detected by FISH using the TelC PNA probe. Co-localization (white arrows) of EdU (cyan) and telomeres (red) appears white, whereas that of PML (green) and telomeres (red) appears yellow. (**D**) The dot plot shows the quantitative numbers of APBs or telomeres in individual BJ^T^ cells (n > 150). A bar chart displays the percentages of APB-positive (APB+) cells, defined as those containing more than five APBs. (**E**) The dot plot shows the quantitative numbers of EdU+Telomere foci in individual BJ^T^ cells (n > 150). A bar chart displays the percentage of EdU+Telomere-positive (ATDS+) cells containing more than three EdU+Telomere co-localizing foci. (**F**) Western blot analysis of TRIM28 expression in U2OS and WI38-VA13/2RA cells after 6 days of treatment with TRIM28-targeting siRNAs. (**G**) Telomere-ChIP analysis of U2OS and WI38-VA13/2RA cells 6 days following TRIM28 knockdown to assess enrichment for telomeric DNA with indicated histones and histone modifications (H3=histone 3, H3K9me3=histone H3 lysine 9 trimethylation, H4=histone 4, H4ac=pan histone H4 acetylation). The relative enrichment was calculated after normalization of ChIP DNA signals to the respective input DNA signals (mean ± SD; n = 3 independent experiments). Results from three independent experiments are shown as individual data points. (**H**) Representative images showing EdU incorporation at telomeres in BJ^T^ cells detected by Click chemistry. PML was detected by IF, and telomeres were detected by FISH using the TelC PNA probe in U2OS and WI38-VA13/2RA cells. Co-localization (white arrows) of EdU (cyan) and telomeres (red) appears white, whereas that of PML (green) and telomeres (red) appears yellow. Dot plots show the quantitative numbers of APBs or EdU+Telomere (ATDS) foci in individual U2OS and WI38-VA13/2RA cells (n > 150). (**I**) Representative images of single-stranded C-rich telomeric DNA (ssTeloC) in U2OS and WI38-VA13/2RA cells. The dot plot shows the quantitative numbers of ssTeloC foci in individual cells. ssTeloC was detected by native FISH using the TelG PNA probe. Red lines in the dot plots indicate the median. (C, G) are determined by the unpaired t-test. (D, E, H, I) are determined by Mann-Whitney U test. Statistical significance is indicated as follows: ns (P > 0.05), * (P < 0.05), ** (P < 0.01), *** (P < 0.001), **** (P < 0.0001).

Next, we examined the ALT-promoting function of TRIM28 in ALT cells. To this end, we depleted TRIM28 from U2OS and WI38-VA13/2RA cells by means of siRNAs (Figure 3F) to investigate how TRIM28 is involved in regulating H3K9me3 and ALT activity. Through telomere-ChIP, we detected that telomeric H3K9me3 was abolished upon depleting TRIM28 from the U2OS and WI38-VA13/2RA cells, supporting that TRIM28 promotes telomeric H3K9me3 in ALT cells (Figure 3G). In addition, we examined whether ALT activity is reduced upon TRIM28 depletion. Indeed, knockdown of TRIM28 from U2OS and WI38-VA13/2RA cells reduced their APB formation and ATDS (Figure 3H). We also performed native fluorescence in situ hybridization (FISH) assay to detect single-stranded C-rich telomeric DNA (ssTeloC), representing another hallmark of ALT activity(Frank et al., 2022). We found that TRIM28 knockdown limited ssTeloC levels in U2OS and WI38-VA13/2RA cells (Figure 3I), indicative of reduced ALT activity. Together, these findings uncover a role for TRIM28 in promoting ALT activity and requisite phenotypes in ALT cells.

### Orphan NRs recruit TRIM28 to telomeres

Given that TRIM28 is required for orphan NR-induced chromatin changes and ALT induction, we wondered if orphan NRs mediate the telomeric localization of TRIM28. To test this possibility, we performed ChIP analysis to examine the association of TRIM28 with telomeres in U2OS and WI38-VA13/2RA cells. Our ChIP results reveal that anti-TRIM28 antibodies pulled down telomere DNA, evidencing the telomeric association of TRIM28 (Figure 4B). Furthermore, following simultaneous depletion of COUP-TF2 and TR4, the enrichment of TRIM28 at telomeres of U2OS and WI38-VA13/2RA cells was diminished (Figure 4A, B), supporting that orphan NRs regulate the telomere association of TRIM28 in ALT cells. By combining immunofluorescence (IF) staining of TRIM28 and telomere-FISH in WI38-VA13/2RA cells, we consistently detected a telomeric localization of TRIM28 and then quantified TRIM28 telomeric enrichment by measuring the signal intensity ratio between the telomere region and the nucleus (Figure 4C). TRIM28 signal intensity at telomeres was reduced following COUP-TF2 plus TR4 depletion (Figure 4C). Similarly, TRIM28-IF and telomere-FISH detected a telomeric localization for TRIM28 in BJ^T^ cells expressing COUP-TF2^LBD^-TRF1, but not in those expressing vector or TRF1 alone (Figure 4D, E). These results further indicate that the telomere association of orphan NRs can promote telomeric enrichment of TRIM28. Importantly, telomeric tethering of the COUP-TF2 LBD having a truncated AF2 domain (COUP-TF2^LBDΔAF2^-TRF1) in BJ^T^ cells failed to induce TRIM28 telomeric localization (Figure 4D, E). Since the AF2 domain of orphan NRs is important for recruitment of co-activators (Kruse et al., 2008) and ALT induction(Gaela et al., 2024), this finding supports that the transcriptional regulatory activity of the COUP-TF2 LBD is critical for the telomeric recruitment of TRIM28. Robust telomeric TRIM28 localization was also observed for KRAB-TRF1-expressing cells (Figure 4D, E), consistent with the direct physical interaction between KRAB and TRIM28 reported previously (Stoll et al., 2022), and serving as a positive control for our experiments. Together, these data evidence that orphan NRs promote the localization of TRIM28 to telomeres.

**Figure 4.**
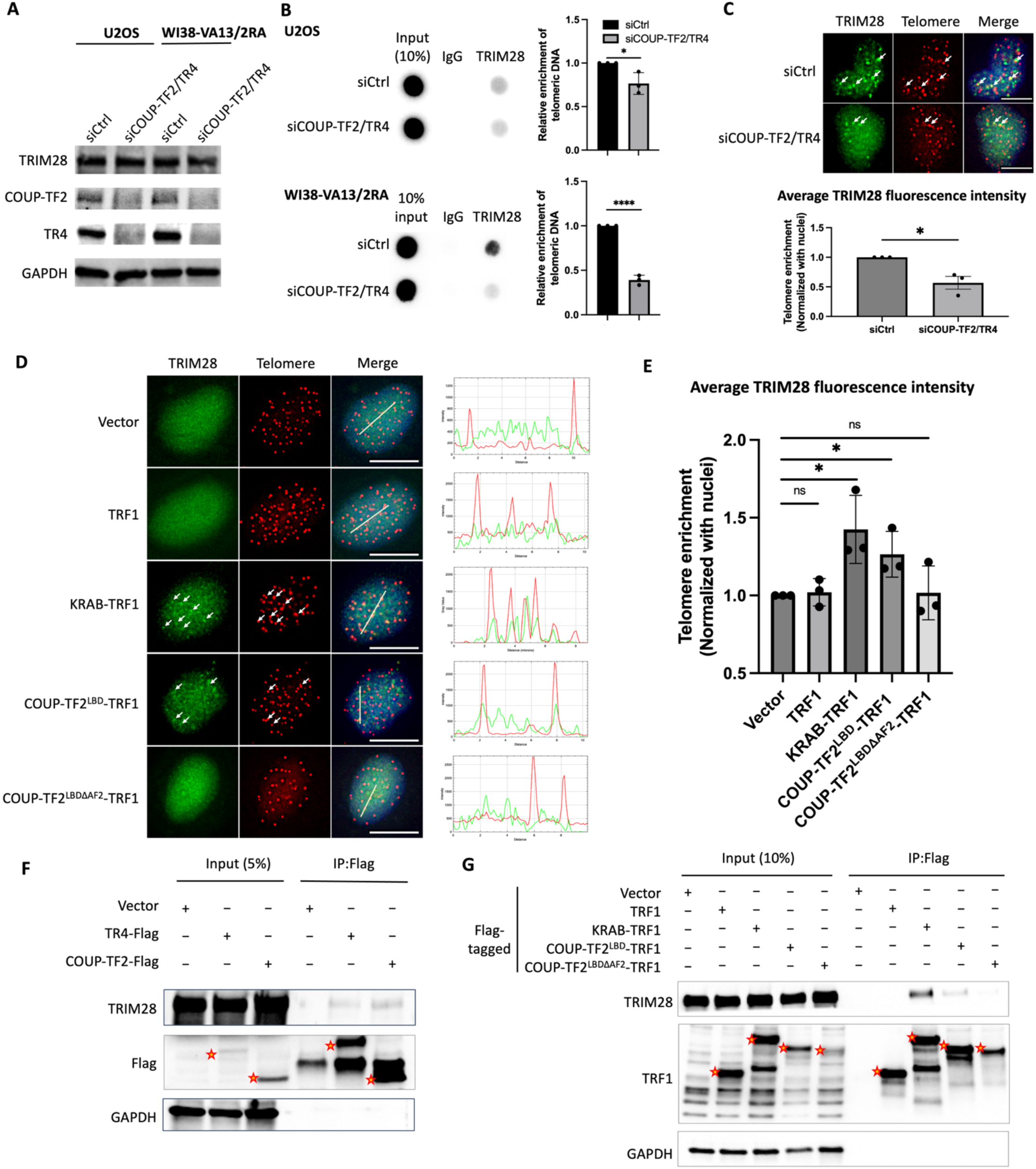
Orphan NRs recruit TRIM28 to telomeres. (**A**) Western blot analysis of TRIM28, COUP-TF2, and TR4 expression in U2OS and WI38-VA13/2RA cells following 6 days of treatment with COUP-TF2- and TR4-targeting siRNAs. (**B**) Telomere-ChIP analysis of BJ^T^ cells 6 days after NR knockdown to quantify enrichment of TRIM28-associated telomeric DNA. The relative enrichment was calculated after normalization of ChIP DNA signals to the respective input DNA signals. Results from three independent experiments are shown as individual data points. (mean ± SD; n = 3 independent experiments). (**C**) Immunofluorescence images depicting TRIM28 localization at telomeres in WI38-VA13/2RA cells after 6 days of NR-targeting siRNA treatment. The bar chart quantifies the average intensity of TRIM28 in the telomeric region normalized to control cells treated with control siRNA, with results from three independent experiments shown as individual data points. (**D**) Representative immunofluorescence images showing TRIM28 localization and telomeres in BJ^T^ cells expressing various plasmids. Intensity plots across the line section are shown beside each image. (**E**) The bar chart quantifies the average TRIM28 fluorescence intensity in the telomeric regions divided by intensity in the nucleus and normalized by the vector expressing cells. Results from three independent experiments are shown as individual data points. (**F**) Co-immunoprecipitation (IP) of TRIM28 with COUP-TF2/TR4. Lysates from WI38-VA13/2RA cells were subjected to IP with TRIM28 and Flag antibodies and analyzed by Western blot. (**G**) Co-immunoprecipitation of TRIM28 with fusion proteins. Lysates from BJ^T^ cells were subjected to IP with TRIM28 and TRF1 antibodies and analyzed by Western blot. (B, C, E) are determined by the unpaired t-test. Statistical significance is indicated as follows: ns (P > 0.05), * (P < 0.05), ** (P < 0.01), *** (P < 0.001), **** (P < 0.0001).

We further investigated whether orphan NRs interact directly with TRIM28 and thereby facilitate TRIM28 telomeric localization. To this end, first, we transiently expressed Flag-tagged COUP-TF2 or TR4 in WI38-VA13/2RA cells for co-immunoprecipitation (Co-IP) using anti-Flag antibodies (Figure 4F). Western blotting revealed that endogenous TRIM28 co-immunoprecipitated with ectopically expressed COUP-TF2 or TR4 in the WI38-VA13/2RA cells (Figure 4F), supporting an association between COUP-TF2/TR4 and TRIM28. We also performed Co-IP with anti-Flag antibodies on BJ^T^ cells stably expressing vector, TRF1, KRAB-TRF1, COUP-TF2^LBD^-TRF1, or COUP-TF2^LBDΔAF2^-TRF1, all tagged with the Flag epitope. The Co-IP results demonstrate that TRIM28 is associated with COUP-TF2^LBD^-TRF1, but not TRF1 alone (Figure 4G). Consistent with its known function, KRAB-TRF1 also co-immunoprecipitated TRIM28 (Figure 4G). Intriguingly, the ability of COUP-TF2^LBD^-TRF1 to interact with TRIM28 and to promote its telomere localization was dependent on the AF2 domain. Together, these findings indicate that orphan NRs may facilitate recruitment of TRIM28 to telomeres through physical interactions.

### The RBCC domain of TRIM28 mediates the COUP-TF2 interaction responsible for telomeric recruitment

To determine the domain(s) of TRIM28 responsible for its telomeric recruitment and ALT-promoting functions, we constructed plasmids to express wild-type TRIM28 protein or mutant variants lacking either the RING Finger-B Box-Coiled Coil (RBCC), PxVxL, or plant homeodomain and bromodomain (PHD/BROMO) domain (Figure 5A). The RBCC domain mediates the interaction between TRIM28 and KRAB domain-containing zinc finger proteins, facilitating TRIM28’s association with specific genomic loci(Wolf et al., 2015). The PxVxL domain is responsible for binding heterochromatin protein 1 (HP1), whereas the PHD/BROMO domain recruits SETDB1 methyltransferase and the NuRD complex, both essential for heterochromatin formation and maintenance(Wolf et al., 2015). We ectopically expressed individual TRIM28 proteins hosting an HA tag in WI38-VA13/2RA cells and then assessed their expression levels and cellular localization by IF-based staining of samples directly fixed with paraformaldehyde. We found that the ectopically-expressed wild-type and mutant TRIM28 proteins primarily localized in the nucleus (Figure 5B), indicating that the RBCC, PxVxL, and PHD/BROMO domains are not required for TRIM28 nuclear localization. To further investigate the chromatin or telomere localization of the TRIM28 proteins, we isolated cytoplasmic and nuclear soluble fractions by incubating the samples with permeabilization buffer containing 0.5% Triton X-100 prior to fixation for IF-telomere FISH (Figure 5C). The results revealed significant retention of wild-type TRIM28 signal in the nucleus, with puncta co-localizing with telomeres (Figure 5c), confirming the association of TRIM28 with chromatin and telomeres in WI38-VA13/2RA cells shown in Figures 4B and 4C. However, deletion of the TRIM28 RBCC domain abolished its nuclear localization and telomeric enrichment (Figure 5C), revealing that the RBCC domain is required for the TRIM28 chromatin association. This result is consistent with the finding previously that TRIM28 associates with chromatin by interacting through its RBCC domain with DNA-binding factors (Wolf et al., 2015). Intriguingly, deletion of the PxVxL domain or PHD/BROMO domain did not affect TRIM28 telomeric enrichment (Figure 5C), so the co-factor recruitment associated with these domains may not be essential for TRIM28’s chromatin or telomeric localization.

**Figure 5.**
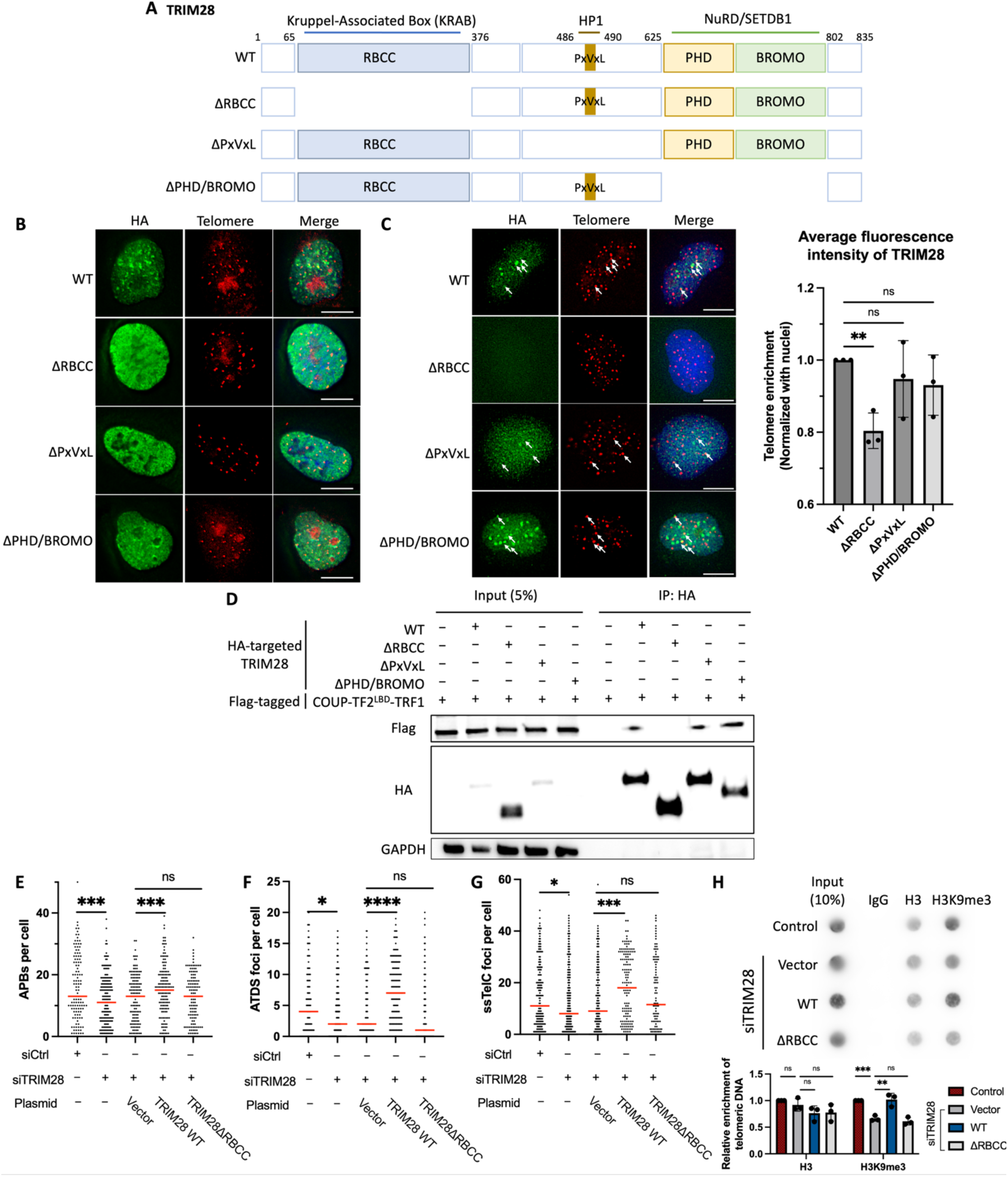
The RBCC domain functions in recruiting TRIM28 to telomeres. (**A**) Schematic representations of wild-type (WT) TRIM28 and its mutant variants. WT TRIM28 comprises an N-terminal RBCC (KRAB binding domain), a central PxVxL (HP1-binding) domain, and a C-terminal PHD/BROMO (NuRD- and SETDB1-interacting) domain. (**B**) Immunofluorescence images showing TRIM28 localization in the nucleus of WI38-VA13/2RA cells following ectopic overexpression of TRIM28 and its mutants without CSK buffer pretreatment. (**C**) Immunofluorescence images showing TRIM28 localization at telomeres in WI38-VA13/2RA cells after ectopic overexpression of TRIM28 and its mutants with CSK buffer pretreatment. The bar chart quantifies the average signal intensity of TRIM28 in the telomeric regions divided by intensity in the nucleus and normalized by the wild-type TRIM28 expressing cells. Results from three independent experiments are shown as individual data points. (**D**) Co-immunoprecipitation of COUP-TF2^LBD^-TRF1 with WT TRIM28 or mutant forms. Lysates from HEK293T cells were subjected to IP with HA and Flag antibodies and analyzed by Western blot. (**E**) The dot plot shows the quantitative numbers of APBs (Telomere+PML) in WI38-VA13/2RA cells after 6 days of TRIM28 knockdown and 2 days of WT TRIM28 or mutant TRIM28 expression (n > 150). (**F**) The dot plot shows the quantitative numbers of EdU+Telomere (ATDS) foci in individual WI38-VA13/2RA cells after 6 days of TRIM28 knockdown and 2 days of WT TRIM28 or mutant TRIM28 expression (n > 150). (**G**) The dot plot shows the quantitative numbers of ssTeloC foci in individual WI38-VA13/2RA cells after 6 days of TRIM28 knockdown and 2 days of WT TRIM28 or mutant TRIM28 expression. ssTeloC was detected by native FISH using the TelG PNA probe (n > 150). (**H**) Telomere-ChIP analysis of WI38-VA13/2RA cells after 6 days of TRIM28 knockdown and 2 days of WT TRIM28 or mutant TRIM28 expression to assess enrichment for telomeric DNA with histone 3 (H3) and histone H3 lysine 9 trimethylation (H3K9me3). The relative enrichment was calculated after normalization of ChIP DNA signals to the respective input DNA signals. Results from three independent experiments are shown as individual data points (mean ± SD; n = 3 independent experiments). (C, H) are determined by an unpaired t-test. (E, F, G) are determined by the Mann-Whitney U test. Statistical significance is indicated as follows: ns (P > 0.05), * (P < 0.05), ** (P < 0.01), *** (P < 0.001), **** (P < 0.0001).

Thus, the RBCC domain is required for TRIM28 to localize at telomeres, indicating that it may also mediate the TRIM28 and COUP-TF2 interaction. To address this possibility, we transiently expressed Flag-tagged COUP-TF2^LBD^-TRF1 with HA-tagged wild-type or mutant TRIM28 in HEK-293T cells for Co-IP using anti-HA antibodies. The Co-IP results revealed that COUP-TF2^LBD^-TRF1 interacts with wild-type TRIM28, as well as with TRIM28 lacking the PxVxL or PHD/BROMO domains, but not with TRIM28 lacking the RBCC domain (Figure 5D), supporting that the RBCC domain is necessary for the interaction between COUP-TF2^LBD^ and TRIM28. Collectively, these findings provide evidence that TRIM28 is recruited to telomeres by COUP-TF2 through an interaction between the RBCC domain of TRIM28 and the LBD of COUP-TF2.

Since the RBCC domain of TRIM28 appears to be critical for recruitment of the protein to telomeres, we anticipated its loss would impair TRIM28’s role in maintaining ALT features and telomeric H3K9me3 levels. To test this hypothesis, we ectopically expressed wild-type TRIM28 or RBCC-deleted TRIM28 in WI38-VA13/2RA cells from which endogenous TRIM28 had been depleted by means of siRNAs (Supplementary Figure 4A), and then assessed the impact on ALT phenotypes. Expression of wild-type TRIM28 restored APB formation, ATDS, ssTeloC, and telomeric H3K9me3 levels in those cells, whereas RBCC-deleted TRIM28 failed to do so (Figure 5E-H). These findings demonstrate that the RBCC domain of TRIM28 is essential for the protein’s function in mediating ALT induction and telomeric chromatin modifications, underscoring its crucial role in facilitating TRIM28’s COUP-TF2 interaction for its telomeric recruitment.

## Discussion

We have demonstrated previously that orphan NRs initiate APB formation and activate ALT pathways when binding to telomeres(Gaela et al., 2024). However, the specific role of chromatin modifications, particularly H3K9me3, in this process had not been defined. Herein, we reveal that targeting the COUP-TF2^LBD^ to telomeres in BJ fibroblasts enhances telomeric H3K9me3 levels (Figure 1D). Conversely, depletion of COUP-TF2 or TR4 in ALT cells (U2OS and WI38-VA13/2RA) reduces telomeric H3K9me3 (Figure 1B). These results establish a direct link between orphan NR-mediated ALT activation and telomeric enrichment of H3K9me3. Mechanistically, we show that orphan NRs recruit TRIM28 to telomeres to promote H3K9me3 and ALT activity (Figure 3B, D, E). This finding is consistent with prior studies reporting that TRIM28 preferentially localizes to ALT telomeres for H3K9me3 enrichment(Czerwińska et al., 2017; Wang et al., 2021). Crucially, we demonstrate here that the interaction between orphan NRs and TRIM28 depends on the RBCC domain of TRIM28 (Figure 4F, 5D). This domain is essential for TRIM28’s recruitment to telomeres, its interaction with COUP-TF2, and its role in facilitating ALT induction and telomeric H3K9me3 activation (Figure 5C-H). Since SETDB1 functions as a cofactor of TRIM28 and is recruited by it to establish H3K9me3 marks (Wang et al., 2021), we propose that orphan NRs recruit TRIM28, which in turn interacts with SETDB1 to promote telomeric H3K9me3. Supporting this hypothesis, we found that SETDB1 depletion significantly reduced COUP-TF2-mediated ALT phenotypes (Supplementary Figure 2C, D), implying a role for SETDB1 in orphan NR-mediated ALT induction. Beyond its role in H3K9me3 activation, TRIM28 interacts with other chromatin regulators, including HP1α/β/γ, HDAC1/2, and DNMT1, suggesting a more extensive role in chromatin remodeling (Czerwińska et al., 2017). These interactions may coordinate additional modifications, such as histone deacetylation or DNA methylation, to create a chromatin environment conducive to ALT maintenance (Figure 6). Further studies are needed to elucidate any additional chromatin-regulatory roles of TRIM28 in orchestrating orphan NR-driven chromatin remodeling and ALT pathway activation at telomeres.

**Figure 6.**
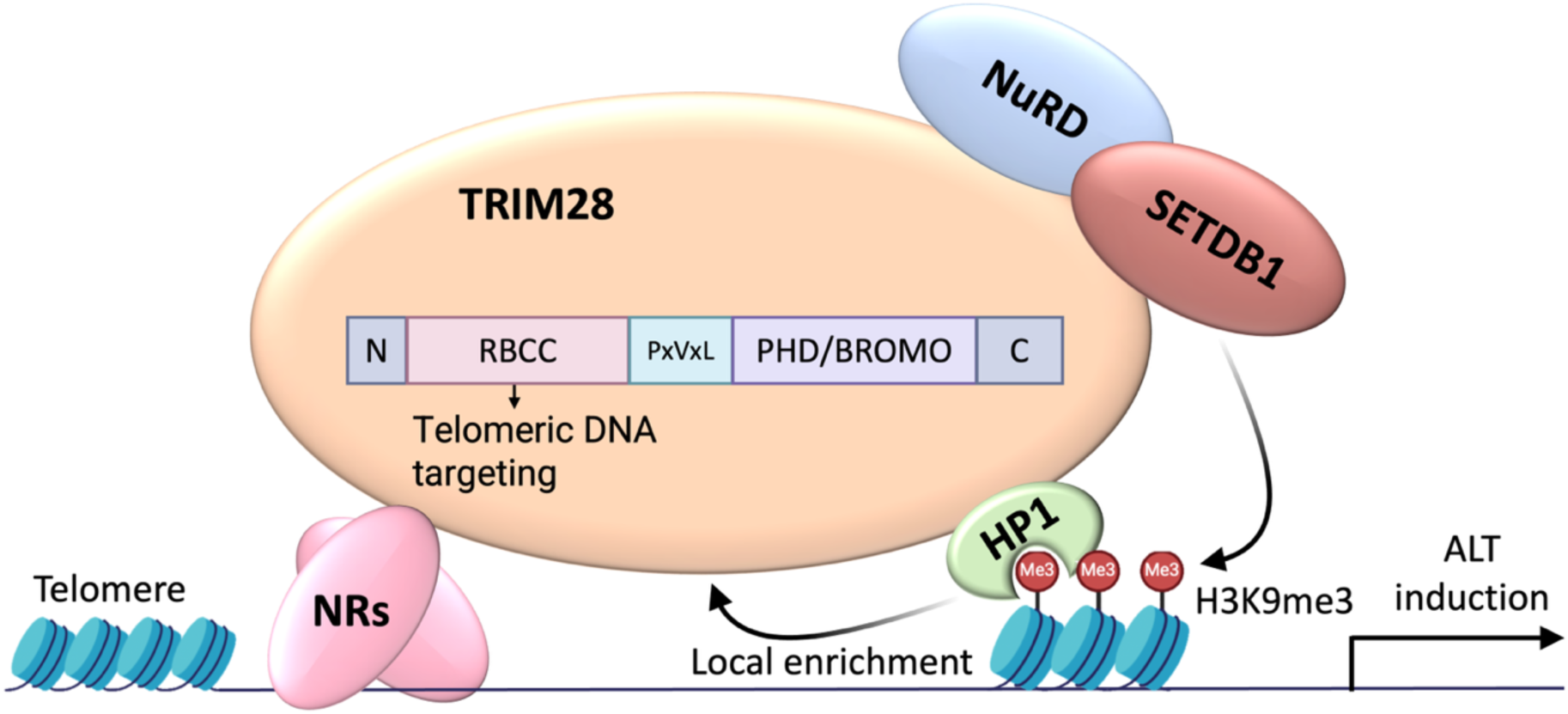
Model depicting how TRIM28 mediates orphan NR-induced ALT activation and telomeric H3K9me3 formation.

H3K9me3, a hallmark of heterochromatin, is significantly enriched at ALT telomeres compared to non-ALT telomeres, defining a distinct feature of the chromatin landscape in ALT cells (Episkopou et al., 2014; Gauchier et al., 2019). While H3K9me3 is known for its role in chromatin compaction, its specific contribution to ALT activation remains incompletely understood. A key function of H3K9me3 is to recruit heterochromatin protein 1 (HP1), a family of proteins integral to heterochromatin formation and maintenance (Bannister et al., 2001; Lachner et al., 2001; Sims et al., 2003). All three HP1 isoforms—HP1α, HP1β, and HP1γ—have been shown to colocalize with APBs (Jiang et al., 2009). Notably, depletion of HP1α or HP1γ from the ALT cell line IIICF-T/B3 leads to significantly reduced APB formation, indicating essential roles for both isoforms in organizing ALT-specific chromatin structures (Jiang et al., 2009). Beyond chromatin association, HP1 proteins mediate liquid–liquid phase separation (LLPS), a critical process in heterochromatin organization(Larson et al., 2017; Tortora et al., 2023). Given that APB formation is triggered by LLPS processes (Banani et al., 2016; Corpet et al., 2020) and requires HP1 (Jiang et al., 2009), we propose that HP1 facilitates APB assembly at ALT telomeres through LLPS mechanisms. This model suggests that the enrichment of H3K9me3 at ALT telomeres promotes APB formation by serving as a scaffold for HP1 recruitment and LLPS-driven chromatin condensation. In addition, HP1 interacts with TRIM28 through its PxVxL motif and plays a pivotal role in stabilizing the TRIM28-containing chromatin complex, which reinforces further H3K9me3 and the establishment of heterochromatin (Czerwińska et al., 2017; Sripathy et al., 2006). Future studies could investigate whether the interaction between TRIM28 and HP1 plays a regulatory role in APB formation and subsequent ALT activation.

In our study, expression of COUP-TF2^LBD^-TRF1 and KRAB-TRF1 in human fibroblasts induced telomeric H3K9me3 and APB formation, demonstrating their ability to alter telomeric chromatin and initiate ALT activity. However, their effectiveness in promoting telomeric DNA synthesis differs significantly. Despite both constructs inducing comparable levels of APB formation (∼60%), KRAB-TRF1 promoted ATDS in less than 8% of the cells, whereas COUP-TF2^LBD^-TRF1 induced ATDS in approximately 20% of the cells (Figure 2C, D, and Figure 3D, E). This disparity may arise from the following possibilities. First, the KRAB-TRF1 fusion protein effectively compacts telomeric chromatin, as indicated by increased histone H3 and H4 occupancy (Figure 2B), which likely impedes access by the recombination machinery and thus limits DNA synthesis. In contrast, COUP-TF2^LBD^-TRF1 leaves histone occupancy unchanged (Figure 1D), maintaining an accessible chromatin state that supports telomeric DNA synthesis.

Second, ATRX, an ALT suppressor, is recruited to telomeres in an H3K9me3-dependent manner(Gauchier et al., 2019). Since KRAB-TRF1 induces greater enrichment of telomeric H3K9me3 than COUP-TF2^LBD^-TRF1 (Figure 1D, 3B), it may also recruit more ATRX, thereby more effectively inhibiting ALT activity. Third, since COUP-TF2 is a transcription factor, it may recruit additional co-activators (Kruse et al., 2008; Xu et al., 2019b) to promote ALT activity. Together, these findings emphasize that though moderate telomeric H3K9me3 enrichment is positively associated with ALT activation, excessive H3K9me3 enrichment or chromatin compaction can hinder telomeric DNA synthesis. This scenario highlights the importance of considering multiple chromatin features, such as histone modifications, chromatin accessibility, and factor recruitment, to understand chromatin’s role in ALT regulation rather than relying on a single characteristic.

Orphan NRs have been shown to promote the recruitment of the ZNF827-NuRD complex to ALT telomeres(Conomos et al., 2014); however, the molecular mechanism remains unclear. TRIM28 interacts with KRAB-domain zinc-finger proteins(Czerwińska et al., 2017). Thus, it is plausible that TRIM28 may act as a protein hub by binding to both orphan NRs and ZNF827, thereby bridging the telomeric recruitment of the NuRD complex and other chromatin remodeling factors for ALT chromatin regulation. Future investigations should examine whether TRIM28 is required for orphan NR-mediated targeting of ZNF827-NuRD complex to telomeres. Moreover, additional zinc-finger proteins, such as ZNF524 and ZBTB40 (BTB/POZ-domain zinc-finger protein), have been implicated in ALT telomere (Braun et al., 2023; Zhou et al., 2023). Thus, it would be valuable to determine if TRIM28 similarly regulates telomeric recruitment of ZNF524 and ZBTB40, potentially uncovering broader mechanisms underlying ALT-dependent telomere maintenance.

## Materials and methods

### Cell lines

BJ^T^ cells, derived from BJ normal fibroblasts stably expressing hTERT, and the human osteosarcoma U2OS and embryonic fibroblast HEK-293T cells (all obtained from ATCC), were cultured in Dulbecco’s Modified Eagle’s Medium (DMEM) supplemented with 10% fetal bovine serum (FBS) and 0.5% penicillin/streptomycin. SV40-transformed WI38/VA13-2RA fibroblasts (also from ATCC) were cultured in Minimum Essential Medium (MEM) containing 10% FBS and 0.5% penicillin/streptomycin. All cell lines were maintained at 37 °C in a humidified atmosphere with 5% CO₂.

Retroviral transduction was used to generate cell lines stably expressing specific proteins. BJ^T^ cells were transduced with Flag-tagged constructs, including vector control, TRF1, KRAB-TRF1, VP64-TRF1, COUP-TF2^LBD^-TRF1, COUP-TF2^LBDΔAF2^-TRF1, COUP-TF2, and TR4. HA-tagged constructs of full-length TRIM28 (amino acids 1–835), TRIM28ΔRBCC (lacking amino acids 65– 376), TRIM28ΔPxVxL (lacking amino acids 486–490), and TRIM28ΔPHD/BROMO (lacking amino acids 625–802) were stably expressed in WI38/VA13-2RA fibroblasts. The TRIM28 plasmid was a gift from Ching-Jin Chang(Chang et al., 2023)

### RNA interference

The siRNAs used in this study were On-TARGETplus reagents (Dharmacon) containing individual or a mixture of four siRNA duplexes generated by the SMARTselection algorithm, with chemical modifications to eliminate off-target effects. Cells were transfected with 25 nM siRNAs using Lipofectamine RNAiMAX (Invitrogen) following the manufacturer’s instructions. Four siRNAs for each of COUP-TF2, TR4, SETDB1, HP1 α/β/γ, HDAC1/2, DNMT1, and TRIM28 were employed. After transfection, cells underwent two rounds of siRNA treatment over a six-day incubation period before being subjected to subsequent analyses.

### Protein extraction and Western blotting

Whole-cell extracts were prepared using 1× SDS sample buffer containing 62.5 mM Tris-HCl (pH 6.8), 10% (v/v) glycerol, 2% (w/v) SDS, 0.01% (w/v) bromophenol blue, and 10% (v/v) β-mercaptoethanol. The extracts were denatured by heating at 90 °C for 10 minutes. Denatured protein samples were resolved on SDS-PAGE gels and electrophoretically transferred to nitrocellulose membranes. The membranes were blocked for 1 hour at 25 °C in Tris-buffered saline (pH 8.0) containing 5% (w/v) dry milk and 0.1% (v/v) Tween-20. Subsequently, membranes were incubated overnight at 4 °C with primary antibodies. The primary antibodies used included anti-COUP-TF2 (PP-H7147-00, R&D Systems), anti-TR4 (PP-H0107B-00, R&D Systems), anti-Flag (SI-F3165, Sigma-Aldrich), anti-HP1 (sc-515341, Santa Cruz), anti-TRIM28 (15202-1-AP, Proteintech), anti-TRF1 (90549, GeneTex), anti-HA (26183, ThermoFisher), and anti-GAPDH (HRP-60004, Proteintech). After washing, the membranes were incubated with horseradish peroxidase (HRP)-conjugated secondary antibodies. Protein bands were detected using SuperSignal West Femto Maximum Sensitivity Substrate (Thermo Fisher Scientific) and visualized via chemiluminescence.

### Detection of telomeric repeat-containing RNA (TERRA)

To detect TERRA expression, total RNA was isolated from BJ^T^ cells using a RNASpin Mini kit (Cytiva) according to the manufacturer’s instructions. Multiple DNase treatments were performed to remove residual genomic DNA, ensuring high RNA purity. Complementary DNA (cDNA) synthesis was carried out using SuperScript III Reverse Transcriptase (Invitrogen) with a TERRA-specific primer (5’-(CCCTAA)_5_-3’) and a housekeeping gene-specific primer. The RNA isolation, DNase treatment, and cDNA synthesis procedures were adapted from an established protocol described by Feretzaki & Lingner (2017).

Quantitative real-time PCR (qPCR) was performed using the Applied Biosystems 7500 Fast Real-Time PCR system (Life Technologies) with a reaction mixture containing 1× SYBR Green Select Master Mix (Life Technologies), cDNA, and specific primer pairs. Primers targeting subtelomeric regions of human chromosomes were designed based on the protocol of Feretzaki & Lingner (2017) and validated using the T2T CHM13v2.0/hs1 genome dataset from Genome Browser (https://hgdownload.soe.ucsc.edu/goldenPath/hs1/bigZips/) to ensure specificity and accuracy. The primer sequences used were 5’-ATGCACACATGACACACACTAAA-3’ (forward) and 5’-TACCCGAACCTGAACCCTAA-3’ (reverse) for the q-arm of chromosome 10, and 5’-GCAAATGCAGCAGTCCTAATG-3’ (forward) and 5’-GACCCTGACCCTAACCCTAA-3’ (reverse) for the q-arm of chromosome 15(Feretzaki & Lingner, 2017).

### Real-time RT-PCR

Total cellular RNA was extracted using an RNASpin Mini kit (Cytiva) following the manufacturer’s instructions. RNA samples were reverse-transcribed into cDNA using iScript Reverse Transcriptase Supermix (Bio-Rad). Real-time PCR analyses were conducted on an Applied Biosystems 7500 Fast Real-Time PCR system (Life Technologies) using a reaction mixture containing 1× SYBR Green Select Master Mix (Life Technologies), cDNA, and specific primer pairs. The primers used for qPCR were: HDAC1: 5ʹ-GGTCCAAATGCAGGCGATTCCT-3ʹ and 5ʹ-TCGGAGAACTCTTCCTCACAGG-3ʹ; HDAC2: 5ʹ-CTCATGCACCTGGTGTCCAGAT-3ʹ and 5ʹ- GCTATCCGCTTGTCTGATGCTC-3ʹ; DNMT1: 5ʹ-GATTTGTCCTTGGAGAACGGTG-3ʹ and 5ʹ- TGAGATGTGATGGTGGTTTGCC-3ʹ; SETDB1: 5ʹ-GCCTACAGCAAGGAACGTATCC-3ʹ and 5ʹ- GTTGATGGCAGGCACACTTGGA-3ʹ; HP1a: 5ʹ-CAATTTCTCAAACAGTGCCGA-3ʹ and 5ʹ- CACCACAGGAATCTGTTGCC-3ʹ; TRIM28: 5ʹ-CAAGATTGTGGCAGAGCGTCCT-3ʹ and 5ʹ- CATAGCCTTCCTGCACCTCCAT-3ʹ; and GAPDH: 5ʹ-GGATTTGGTCGTATTGGG -3ʹ and 5ʹ-GGAAGATGGTGATGGGATT -3ʹ. Cycle threshold (Ct) values were normalized to GAPDH, which was used as an internal control. Relative gene expression levels were calculated using the comparative Ct (ΔΔCt) method.

### Detection of telomeric DNA synthesis

To synchronize cells in the G2 phase, a 21-hour treatment with 2 mM thymidine was applied, followed by a 4-hour release into fresh medium. Subsequently, cells were exposed to 15 mM CDK1 inhibitor for 12 hours. To detect DNA synthesis, cells were incubated with 20 mM EdU for 3 hours, after which EdU incorporation was visualized using a Click-iT reaction with fluorescently-labeled picolyl azide (Invitrogen).

### Immunofluorescence (IF) and fluorescence in situ hybridization (FISH)

Cells grown on coverslips within 12-well plates were fixed using 4% paraformaldehyde for 10 minutes. Permeabilization, performed optionally to remove nuclear soluble fractions, involved treatment with 0.5% Triton X-100 in PBS for 5 minutes. The cells were then blocked in 2% FBS diluted in PBS for 30 minutes and incubated with anti-Flag or anti-PML primary antibodies for 1 hour. Then, Alexa 488-conjugated secondary antibodies diluted in 2% FBS were applied for another hour. After immunostaining, the cells were re-fixed with 4% paraformaldehyde for 10 minutes and dehydrated through a series of ethanol treatments (70%, 95%, and 100%) for 5 minutes each. Once air-dried, the coverslips were subjected to hybridization using a mixture containing 10 mM Tris–HCl (pH 7.5), 70% formamide, 10% blocking reagent (Roche), and 0.25 μM TMR-conjugated (CCCTAA)₃ telomere PNA probe. Denaturation was conducted at 80 °C for 3 minutes on a heat plate, followed by overnight hybridization at room temperature. Post-hybridization washes consisted of two rinses with wash buffer A (70% formamide, 10 mM Tris–HCl, pH 7.5) and three with wash buffer B (10 mM Tris–HCl, pH 7.5, 4 M NaCl, 20% Tween-20). The nuclei were stained with DAPI (Life Technology), followed by a second ethanol dehydration series and air-drying of the coverslips. Finally, the coverslips were mounted using ProLong Gold Antifade Mountant (Life Technology).

### Telomere-ChIP

Cells were cultured in 15-cm dishes and cross-linked with 1% formaldehyde at room temperature for 10 minutes. The reaction was quenched by washing the cells twice with PBS containing 200 mM glycine (PBS/Glycine). The cells were scraped into 5 mL of PBS/Glycine, centrifuged, and washed with PBS. The cell pellet was resuspended in 1 mL of Swelling Buffer, incubated on ice for 10 minutes, and centrifuged at 4000 rpm. The pellet was resuspended in 600 μL of sonication buffer and subjected to 40 cycles of sonication (30 seconds on, 30 seconds off) using a Bioruptor. The lysate was clarified by centrifugation at maximum speed for 15 minutes, and the supernatant was divided into five 120 μL aliquots.

For immunoprecipitation (IP), the supernatant was diluted four-fold with IP dilution buffer (16.7 mM HEPES, 150 mM NaCl, 1 mM EDTA, 0.1% sodium deoxycholate, 0.1% SDS, 1% Triton X-100, and protease inhibitors) and incubated overnight at 4 °C with 5 μg of antibody and 20 μg of bacterial DNA, reserving 5–10% of the IP mix as input. The following day, 40 μL of Protein G agarose preincubated with 5 μg of bacterial DNA and 30 μg of BSA was added to each sample and incubated for 30 minutes at 4 °C. The beads were washed sequentially with Wash A, Wash B, Wash C, and TE buffers. Chromatin was eluted in three 150 μL aliquots of Elution Buffer, and reverse cross-linking was performed by adding 200 mM NaCl and incubating the samples at 65 °C for 4 hours. The antibodies used included anti-Histone 3 (ab1791, Abcam), anti-Histone H3 (acetyl K9) (ab4441, Abcam), anti-Histone H4 (14149S, Cell Signaling), anti-acetyl-Histone H4 (06-866, Merck), and anti-TRIM28 (15202-1-AP, Proteintech).

The eluted DNA was treated with 10 μL of 0.5 M EDTA, 20 μL of 1 M Tris-HCl (pH 6.5), and 5 μg of RNase A, followed by incubation at 37 °C for 1 hour. Subsequently, 40 μg of Proteinase K was added, and the samples were incubated at 50 °C for 1 hour. DNA was purified using a Qiagen PCR purification kit. The purified DNA (50 μL) was mixed with 50 μL of 0.46 M NaOH, heated at 95 °C for 5 minutes, and cooled on ice before neutralization.

Dot blotting was performed on Hybond XL membranes (GE Healthcare Life Sciences) following the manufacturer’s protocol. Membranes were air-dried for 5 minutes, cross-linked with 240 mJ using a Stratalinker at 254 nm, and prehybridized in Church hybridization buffer (1% BSA, 1 mM EDTA, 500 mM NaHPO_4_ [pH 7.2], and 7% SDS). Hybridization was carried out overnight with a 5’-(CCCTAA)3-3’ DNA oligonucleotide probe end-labeled with [γ-32P]ATP using T4 polynucleotide kinase (Promega). The membranes were washed three times for 5 minutes in 2× SSC, exposed to a phosphor screen, and imaged. Signal quantification was performed using the array analysis function in Fiji, with background subtraction from unused wells. The amount of immunoprecipitated telomeric DNA was normalized for each sample based on input telomeric DNA signal intensities determined from standard curves.

### Immunoprecipitation (IP)

Cells cultured in 10 cm plates were prepared for immunoprecipitation as follows. The supernatant was removed, and the cells were washed with 1 mL of PBS. The cell suspension was transferred to an Eppendorf tube, centrifuged at 4 °C, and the supernatant was discarded. The pellet was washed twice with 1 mL of PBS, with centrifugation performed after each wash. The washed pellet was resuspended in 570 μL of NETN lysis buffer (1 mM EDTA, 40 mM Tris, 100 mM NaCl, 0.5% V/V NP-40) containing protease inhibitors (ROC-04693132001, Roche), vortexed, and centrifuged at maximum speed for 10 minutes at 4 °C. The resulting supernatant was transferred to a new tube, centrifuged again, and 930 μL of the cleared lysate was used for immunoprecipitation, with 50–25 μL reserved as an input control. For immunoprecipitation, 30 μL of anti-Flag agarose beads (A2220, Sigma-Aldrich) were added to each sample, and the mixture was incubated at 4 °C for 30 minutes with gentle rotation. The beads were washed with NETN buffer and resuspended in 70 μL of 2X SDS loading buffer containing 5% β-mercaptoethanol. The samples were incubated at room temperature for 10 minutes, centrifuged, and the supernatant containing the immunoprecipitated proteins was collected for analysis.

### Image acquisition and quantification

Fluorescence 3D images were acquired using a GE Healthcare DeltaVision Deconvolution microscope with 0.2 μm spacing for a total of 5 μm. The images were processed with software version 5.5.1 for deconvolution and analyzed using ImageJ/FIJI for quantification. Initially, a maximum intensity projection was applied to convert the Z-stack into a 2D image. Background subtraction was performed using the ’tophat’ filter. To identify foci, we applied Ilastik, a tool trained with an artificial intelligence (AI) model (Berg et al., 2019).

### Statistical analyses

Data visualization, including tables and graphs, was performed using GraphPad Prism 9 and Microsoft Excel. For imaging-based cell analysis, each experiment was independently repeated three times, with a minimum of 50 cells analyzed per replicate, yielding a total of approximately 150–300 cells per experimental condition. In scatter plots, each point represents a quantified value per cell, with red lines denoting either the mean or median for inter-sample comparison. Statistical analyses were conducted using two-tailed Mann– Whitney U tests or unpaired t-tests, depending on the data distribution and experimental design.

## Acknowledgements

We thank L.-R. You of the Institute of Biochemistry and Molecular Biology at National Yang Ming University, S.- J. Chou of the Institute of Cellular and Organismic Biology at Academia Sinica, and C.-J. Chang of the Institute of Biological Chemistry at Academia Sinica for providing reagents, T.-E. Chen for technical help, and S.-P. Lee, Y.-H. Liou and C.-Y. Chang of the Imaging Core in the Institute of Molecular Biology at Academia Sinica for help with data collection.

## Author contributions

C.-T.T., H.-Y.H., V.M.G., and L.-Y.C. designed the study; C.-T.T. performed most of the experiments, imaging, and data analysis, with assistance from H.-Y.H. and Y.-C.H.; C.-T.T., H.- Y.H., and V.M.G. developed the protocol for image analysis; C.-T.T., V.M.G., and L.-Y.C. wrote the manuscript.

## Funding

Research in the laboratory of L.-Y. Chen was supported by grants from Academia Sinica [AS-GCP-113-L02]. Funding for open access charge: Academia Sinica.

## Conflict of interest statement

None declared.

---

**Supplementary Figure 1.**
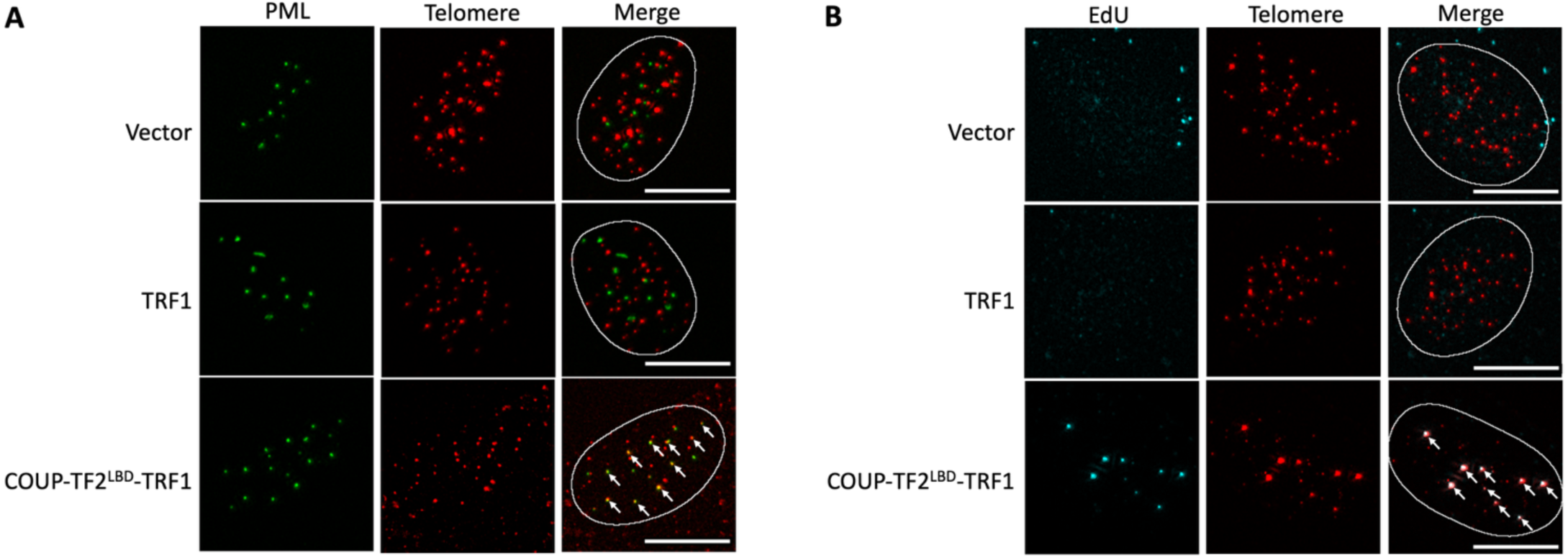
Characterization of COUP-TF2^LBD^-TRF1 in promoting APB formation and ATDS. (a) Representative images showing EdU incorporation at telomeres in BJ^T^ cells detected by Click chemistry. PML was detected by IF, and telomeres were detected by FISH using the TelC PNA probe. Co-localization of PML (green) and telomeres (red) appears yellow. The white outlines encompass DAPI segmentation. White arrows indicate APBs. (b) Representative images showing EdU incorporation at telomeres in BJ^T^ cells expressing vector, TRF1 alone, or COUP-TF2^LBD^-TRF1. IF detects EdU, and telomeres are visualized by FISH using the TelC PNA probe. Co-localization of EdU (cyan) and telomeres (red) appears white. The white outlines encompass DAPI segmentation.

**Supplementary Figure 2.**
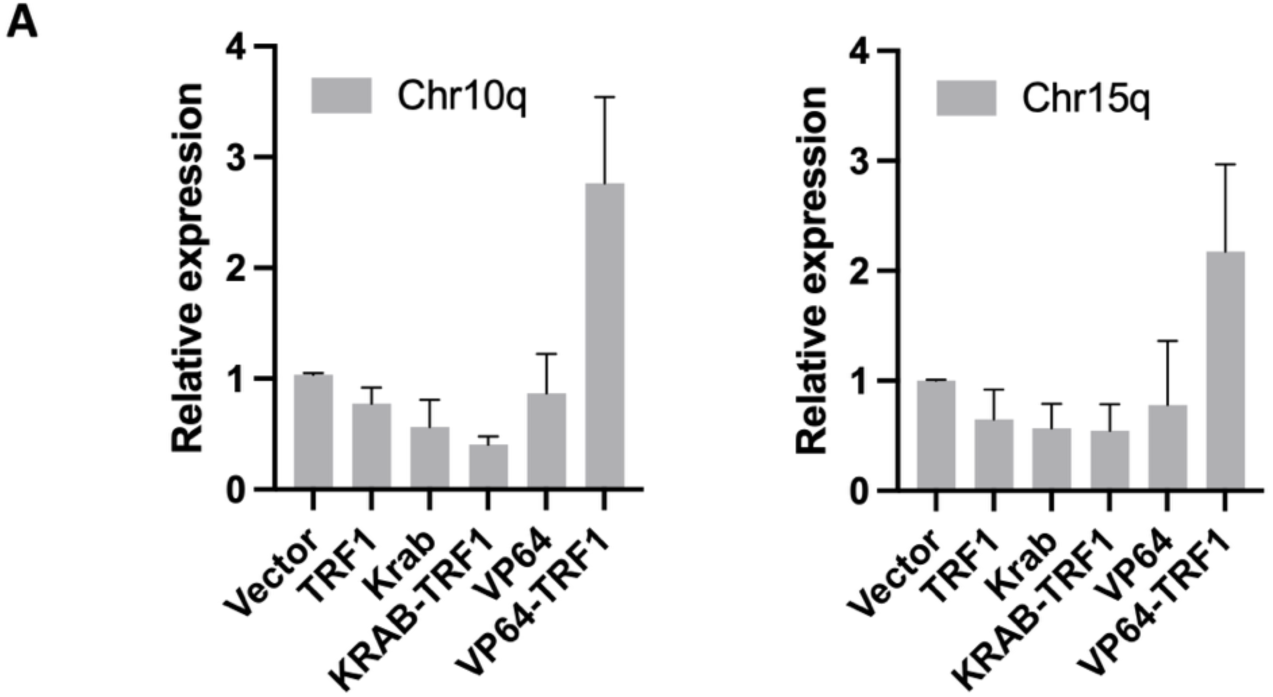
Quantification of TERRA levels at 10q and 15q subtelomeres. (**A**) Quantification of TERRA expression levels in TRF1. KRAB, KRAB-TRF1, VP64, VP64-TRF1 BJ^T^ cells normalized to Flag BJ^T^ cells, as determined by qPCR of RNA samples. The TERRA levels with different subtelomeric sequences from the chromosome (Chr) 10q or 15q ends were measured and normalized to GAPDH. Your Method and panel indicate chromosome 15. (mean ± SEM; n = 5 independent experiments)

**Supplementary Figure 3.**
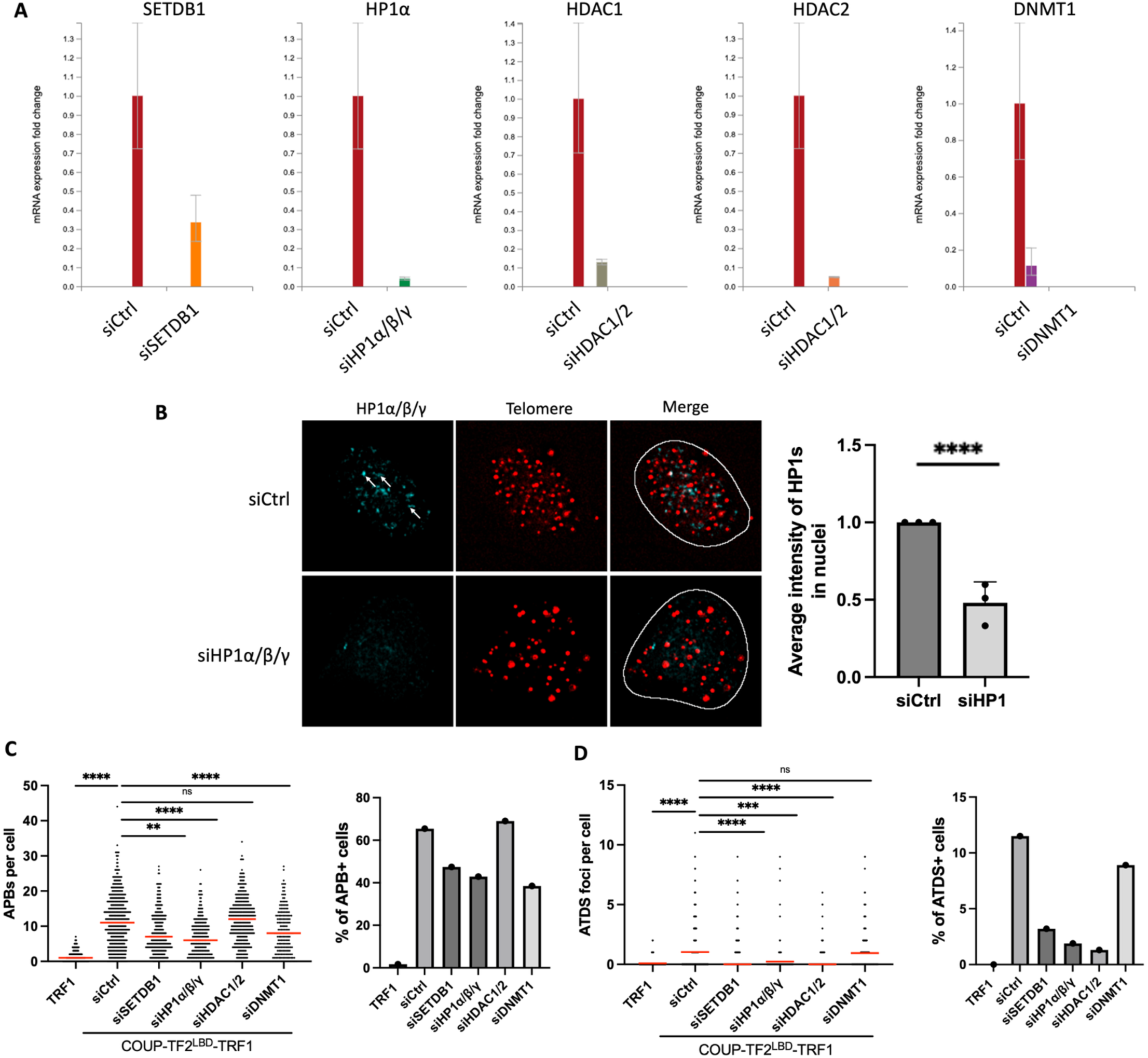
Functional analysis of siRNA-mediated knockdown of heterochromatin-associated proteins in BJ^T^ cells expressing COUP-TF2^LBD^-TRF1. (**A**) qPCR and (**B**) IF data showing the gene silencing efficiency of siRNA sequences targeting specific RNA in BJ^T^ cells expressing COUP-TF2^LBD^-TRF1. The bar chart indicates the average HP1 signal intensity per nucleus. Each dot indicates one independent experiment (mean ± SD; n = 3 independent experiments). (**C**) The dot plots show the quantitative numbers of APBs in TRF1 or COUP-TF2^LBD^-TRF1 BJ^T^ cells treated with specific siRNAs targeting heterochromatin-associated proteins. (**D**) The dot plots show the quantitative numbers of EdU+ APB foci in TRF1 or COUP-TF2^LBD^-TRF1 BJ^T^ cells treated with specific siRNAs targeting heterochromatin-associated proteins. Red lines in the dot plots indicate the median. (B) Statistical significance is indicated as follows: ns (P > 0.05), * (P < 0.05), as determined by unpaired t-test. (C, D) Statistical significance is denoted as follows: ns (P > 0.05), * (P < 0.05), ** (P < 0.01), *** (P < 0.001), **** (P < 0.0001), as determined by the Mann-Whitney U test. All knockdown experiments involved treating cells with specific siRNAs for 6 days.

**Supplementary Figure 4.**
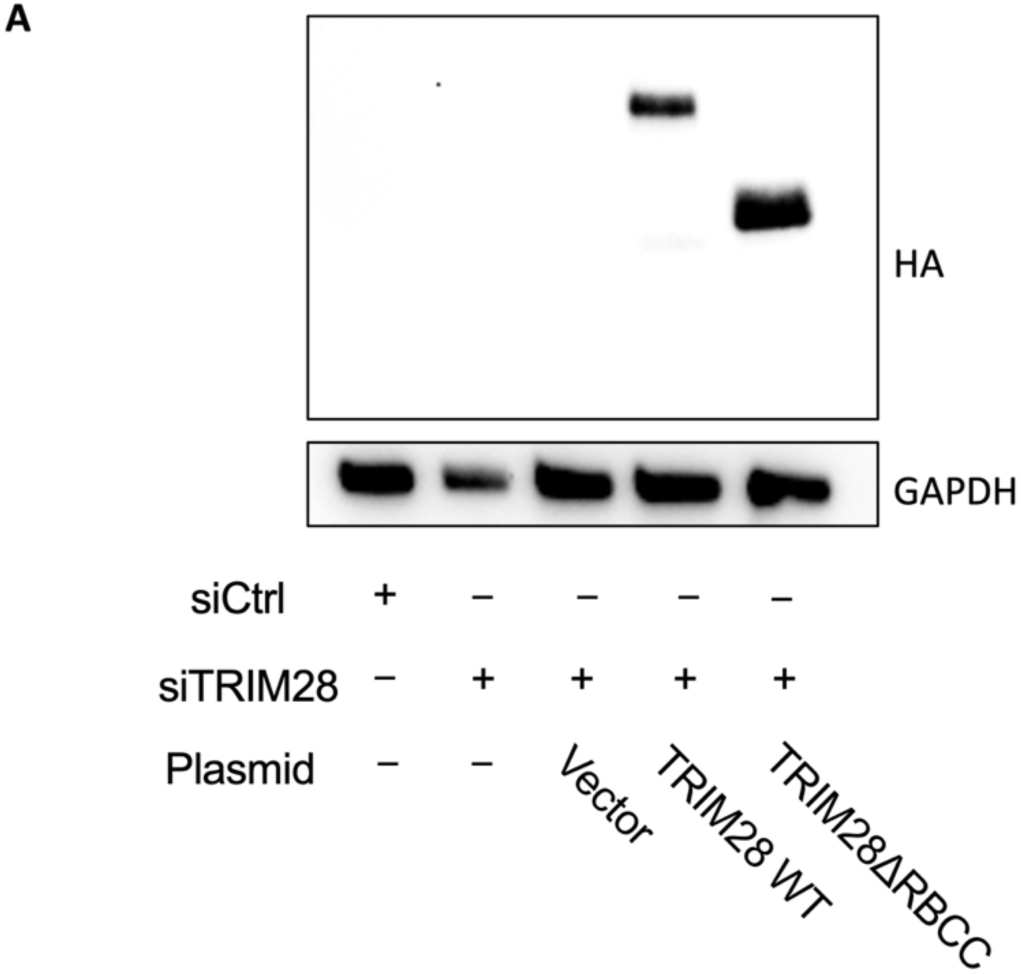
Western blot analysis of mutant TRIM28 expression in TRIM28 knockdown WI38-VA13/2RA cells. (A) Western blot analysis of WI38-VA13/2RA cells after 6 days of TRIM28 knockdown and 2 days of WT TRIM28 or mutant TRIM28 expression, confirming expression of the constructs using an anti-HA antibody.

## Notes

### Competing Interest Statement

The authors have declared no competing interest.

